# Systematic computer-aided disulfide design as a general strategy to stabilize prefusion class I fusion proteins

**DOI:** 10.1101/2024.02.29.582784

**Authors:** Karen J. Gonzalez, Kevin Yim, Jorge Blanco, Marina S. Boukhvalova, Eva-Maria Strauch

## Abstract

Numerous enveloped viruses, such as coronaviruses, influenza, and respiratory syncytial virus (RSV), utilize class I fusion proteins for cell entry. During this process, the proteins transition from a prefusion to a postfusion state, undergoing substantial and irreversible conformational changes. The prefusion conformation has repeatedly shown significant potential in vaccine development. However, the instability of this state poses challenges for its practical application in vaccines. While non-native disulfides have been effective in maintaining the prefusion structure, identifying stabilizing disulfide bonds remains an intricated task. Here, we present a general computational approach to systematically identify prefusion-stabilizing disulfides. Our method assesses the geometric constraints of disulfide bonds and introduces a ranking system to estimate their potential in stabilizing the prefusion conformation. We found, for the RSV F protein, that disulfides restricting the initial stages of the conformational switch can offer higher stability to the prefusion state than those preventing unfolding at a later stage. The implementation of our algorithm on the RSV F protein led to the discovery of prefusion-stabilizing disulfides, providing evidence that supports our hypothesis. Furthermore, the evaluation of our top design as a vaccine candidate in a cotton rat model demonstrated robust protection against RSV infection, highlighting the potential of our approach for vaccine development.

**Significance statement:** Class I fusion proteins, pivotal for cellular entry, constitute primary targets for neutralizing antibodies across a wide range of highly pathogenic viruses. Notably, the prefusion conformation of these proteins has demonstrated to be a robust immunogen, presenting a promising avenue for vaccine development. Despite of this, the inherent structural instability of this conformation poses a substantial hurdle for vaccine formulations. To overcome this limitation and to optimize these proteins as vaccines candidates, we developed an automated computational approach to enhance the structural stability of the prefusion conformation without compromising its immunogenic properties. Given the clinical significance of class I fusion proteins, our computational tool offers a mechanism to accelerate the development of preventive measures against a multitude of viral infections.

## Introduction

Since the emergence of the severe acute respiratory syndrome coronavirus 2 (SARS-CoV-2) and subsequent development of COVID-19 vaccines, it has become apparent that prefusion class I fusion proteins are potent vaccine candidates. This potential is not unique to SARS-CoV-2 and has also been illustrated in other pathogenic viruses, such as Hendra and Nipah (1, 2), parainfluenza virus types 1-4 (3), influenza (4), human metapneumovirus (hMPV) (5), and the respiratory syncytial virus (RSV) (6–8). Despite of this, the intrinsic structural instability of these proteins poses a significant challenge when integrating them into vaccine formulations. As part of their function as membrane-fusion mediators, class I fusion proteins can irreversibly refold from their metastable prefusion state to a highly stable postfusion state (9). This spontaneous transition is a major hurdle in vaccine development, as potent immunogenic epitopes are often hidden in the protein’s most stable structure, the postfusion state (10, 11). Consequently, stabilizing the prefusion conformation has been a continuous pursuit for maintaining the protein’s immunogenicity and leveraging its potential as a vaccine component.

Understanding the conformational rearrangements of class I fusion proteins was largely possible thanks to the crystallization of the influenza virus hemagglutinin protein (12–14) and paramyxoviruses fusion proteins (15–18). As a common fusion mechanism, class I fusion proteins are initially maturated by proteolytic cleavage, where a hydrophobic fusion peptide is left unconstrained to connect with the host cell (19, 20). Following its activation, the fusion-competent yet fragile prefusion state undergoes conformational changes, facilitating the fusion peptide’s interaction with the target membrane (12, 21, 22). As this interaction progresses, the protein’s C-terminal relocates, leading to the assembly of a highly stable six-helix bundle, the postfusion state. This structural transition ultimately enables the fusion of the viral and cellular membranes (23–25).

The structural analysis of the fusion mechanism has guided the prefusion stabilization of various class I fusion proteins by targeting regions prone for refolding (26). Historically, this goal has been achieved through an arduous manual exploration of the protein structure and extensive testing of protein mutants (1, 3, 26–31). We had previously pioneered a computational approach that significantly streamlined this demanding process, requiring only a few designs to identify stable prefusion versions (32). However, while our algorithm enabled the semi-automated introduction of stabilizing mutations using established molecular strategies, such as cavity filling, elimination of buried polar residues, and optimization of polar and electrostatic interactions, it did not allow for the incorporation of disulfide bonds.

To complement our original stabilization strategy, we have developed a general computational approach to systematically identify prefusion-stabilizing disulfides. Our method traces potential disulfides based on allowed distance and geometry, similar to other software (33–35). However, while correct geometry indicates the likelihood of bond formation, it may not directly correlate with stabilizing effects (33), (36). Consequently, we sought to tailor the disulfide design process to class I fusion proteins by considering the conformational dynamics of the protein during the selection of prefusion-stabilizing bonds. We hypothesize that disulfides restricting the initial stages of the conformational switch are more impactful at increasing the stability of the prefusion state than those located on regions unfolding last. We applied our novel disulfide-design strategy to the RSV fusion (F) protein, as we had previously optimized a construct, the R-1b protein, using our first methodology. Consequently, we had an ideal test case to assess the cumulative enhancements offered by the integration of both our approaches. By combining our prior approach with this disulfide-design strategy, we have successfully increased the stability of the RSV F protein without compromising its immunogenic properties.

## Results

### Disulfide bond design strategy

Inspired by the efficacy of non-native disulfides in stabilizing the prefusion conformation of class I fusion proteins (1, 3, 7, 29), we have developed a computational approach to automatically identify these covalent bonds. Building upon a previously proposed concept (5), our computational strategy is centered on identifying novel disulfides within regions undergoing significant relocation during the prefusion-to-postfusion transition (Fig. 1.A). Using PyRosetta as the disulfide scanning tool (37, 38), we first identify potential disulfide bonds via the proximity of Cβ atoms. However, recognizing that proteins are inherently dynamic and that resolved structures represent only single instances in time, we perform this analysis across a spectrum of potential conformations derived from structural energy minimization (39–41). This relaxation process is guided by electron density data to prevent the introduction of artifacts due to excessive deviations from the protein’s original configuration.

**Figure 1.**
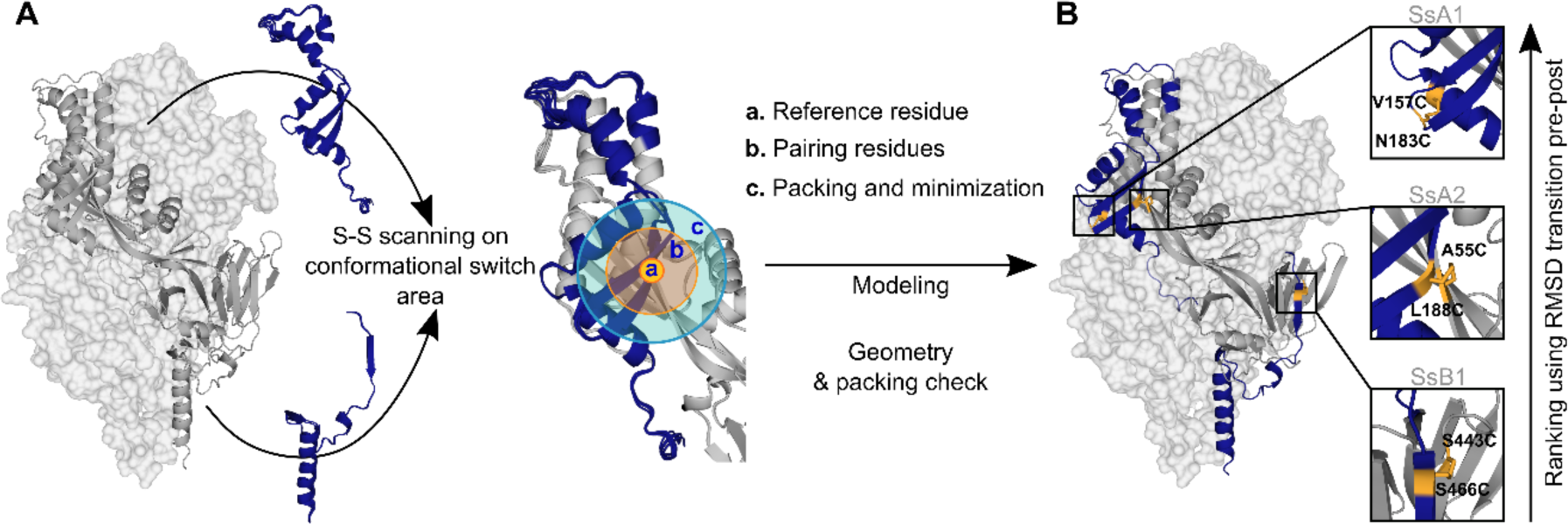
Disulfide design strategy. **(A)** Disulfide scanning focuses on residues located on the protein’s conformational switch area. The protein is analyzed as a dynamic ensemble of structures rather than a static model to identify potential disulfide bonds across various configurations. Each ensemble structure is explored in small sections to streamline computational efforts. These regions are defined by: (a) Reference residue, specifically an amino acid situated in the conformational switch area. (b) Pairing residues, which are adjacent amino acids within a 6Å radius, capable of forming disulfide bonds with the reference residue. (c) Repacking and minimization zone, an area extending 10Å from the pairing residues, where structural repacking and energy optimization occur during the modeling process. **(B)** New potential disulfide bonds identified in R-1b. Following the modeling of potential disulfides, candidates undergo filtration based on correct geometry and non-disruption of protein packing. Ultimately, the newly identified bonds are ranked according to the protein’s conformational dynamic with the assumption that disulfide bonds placed in regions of high root-mean-square-deviation (RMSD) between pre- and postfusion structures would confer higher stability to the prefusion state. The R-1b protein (PDB: 7tn1) is on display with two protomers as light-grey molecular surfaces and one protomer as a dark-grey ribbon. Regions undergoing drastic conformational changes are highlighted in dark blue, and newly designed disulfide bonds are shown in yellow sticks.

In each conformation of the protein ensemble, residues within a 6Å radius are mutated to cysteine, and disulfide bonds are enforced during a focused round of energy minimization and protein repacking (Fig. 1.A). With this focused structural sampling, we not only mitigate computational load but also ensure that the newly introduced disulfide aligns with the specific protein conformation under study. Finally, to increase the probability of successful disulfide formation, we apply a filtering process, assessing potential bonds for Rosetta disulfide energy (dslf_fa13) (42, 43), and geometric adequacy (Fig. 2.A), while ensuring the protein packing is not disrupted.

**Figure 2.**
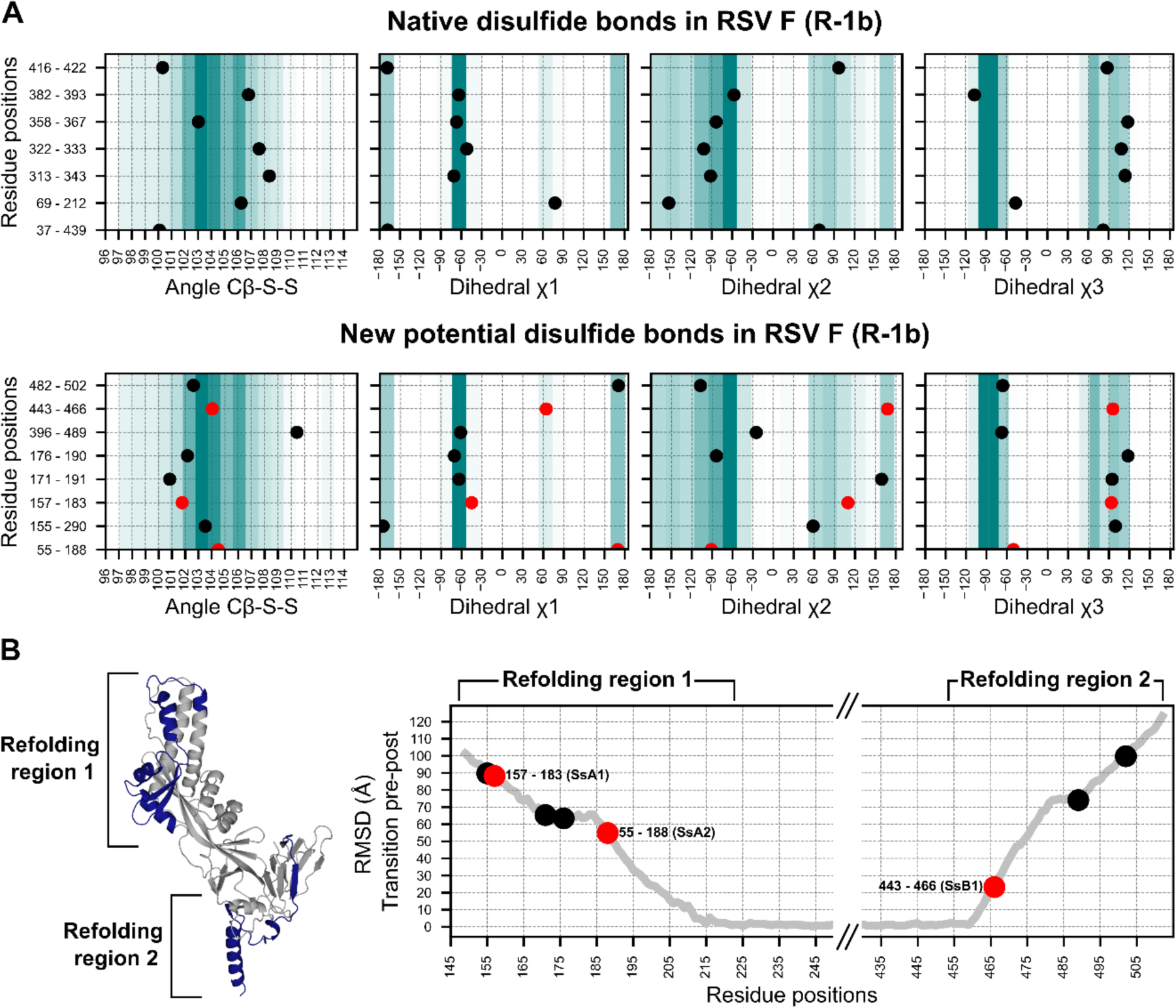
Geometry assessment of the newly identified potential disulfides in the RSV F construct R-1b. **(A)** Geometric parameters used to evaluate the correctness of a disulfide bond. The top panel displays the bond angles sampled by the native disulfides in the R-1b protein while the bottom panel presents the bond angles of the newly identified potential disulfides. The background color scales in each plot represent the frequency of each bond angle, according to a set of 300 high-resolution structures (<1.5Å). Darker colors indicate angles that were more frequent in this reference set, while lighter colors represent angles of low frequency. **(B)** Root-mean-square-deviation (RMSD) mapping of residues forming new potential disulfide bonds. RMSD values were calculated by aligning the prefusion and postfusion structures of the RSV F protein (PDB: 7tn1and 3rrt). The highest RMSD found within the residue pair forming the disulfide bond determined the ranking of the disulfide. Given that conformational changes in the RSV F protein likely occur at distinct times in the refolding regions 1 and 2, the ranking position should be independently analyzed for each region. The R-1b protein (PDB: 7tn1) is displayed, with the conformational switch area highlighted in dark blue. Disulfides chosen for experimental validation are denoted by red circles in all panels.

To evaluate the extent to which disulfide design could complement and enhance our existing computational framework, we applied our methodology to our previously optimized variant of the RSV F, the R-1b protein (32). This implementation led to the identification of 8 potential intrachain disulfides within regions involved in the protein’s conformational switch. Notably, one of these disulfides corresponded to the most effective non-native disulfide previously reported for RSV F, the S155C-S290C disulfide (7) (Fig. 2.A).

With the goal of identifying metrics that could rank disulfides based on their potential stabilizing effects, we analyzed the geometric constraints of disulfides with prior experimental data (7, 26, 44). Interestingly, we observed that a significant number of these previously tested disulfides exhibited appropriate geometric configurations yet failed to enhance protein stability. This observation suggested that other factors beyond geometric parameters might play a role in determining the stabilizing capabilities of disulfide bonds. Consequently, we sought to complement our geometry analysis by exploring the role of the protein’s dynamics in identifying prefusion-stabilizing disulfides. We hypothesized that substitutions restricting the initial conformational changes of the protein would have a stronger impact on the stability of the prefusion state than those preventing the final stages of the rearrangement.

To integrate this concept into our selection process, we assumed that regions displaying higher root-mean-square-deviation (RMSD), when comparing the prefusion and postfusion structures, are more likely to unfold at an earlier stage. As a result, RMSD served as the quantitative metric to rank disulfide candidates according to their potential for prefusion stabilization (Fig. 2.B). Importantly, given that conformational changes in the refolding region 1 (protein head) likely trigger conformational changes in the refolding region 2 (membrane-proximal domain)(30), our ranking system was applied independently to each refolding region.

To validate our design and ranking approaches, we expressed three distinct disulfide variants, each exhibiting well-differentiated RMSD values (Fig. 2.B). The selected disulfides included *V157C-N183C* (design SsA1), located at the beginning of the refolding region 1, *A55C-L188C* (design SsA2), found at the middle of the refolding region 1, and *S443C-S466C* (design SsB1), situated at the beginning of the refolding region 2 (Fig. 1.B and 2.B). Notably, while this manuscript was in preparation, an independent study published experimental validations for the latter two disulfides (45).

### Biochemical characterization of disulfide candidates

All three designs, SsA1, SsA2, and SsB1, were successfully expressed and purified as trimeric proteins, as confirmed by size exclusion chromatography (SEC) (Fig. 3.A). Additionally, the expression yield of these variants showed no significant changes when compared to the parent construct R-1b (Fig. 3.A). As anticipated, the designed proteins exhibited notable improvements in thermal stability compared to R-1b, with melting temperature increments of 2.5°C (SsB1), 5.5°C (SsA2), or 12.5°C (SsA1) (Fig. 3.B). This enhancement in heat stability aligned well with our proposed hypothesis, emphasizing that disulfides restricting the initial stages of protein unfolding can have a more profound impact on the overall stability of the prefusion conformation. Remarkably, the thermal stability of design SsA1 surpassed that of the current RSV vaccine DS-Cav1(7) by approximately 6.5°C (Fig. 3.B). However, it is important to note that these results do not imply that the V157C-N183C disulfide in SsA1 is inherently superior to the S155C-S290C disulfide found in DS-Cav1. Instead, the evident improvement observed in SsA1 likely arises from a synergistic effect resulting from the introduction of the new disulfide and the preexisting mutations in R-1b (32).

**Figure 3.**
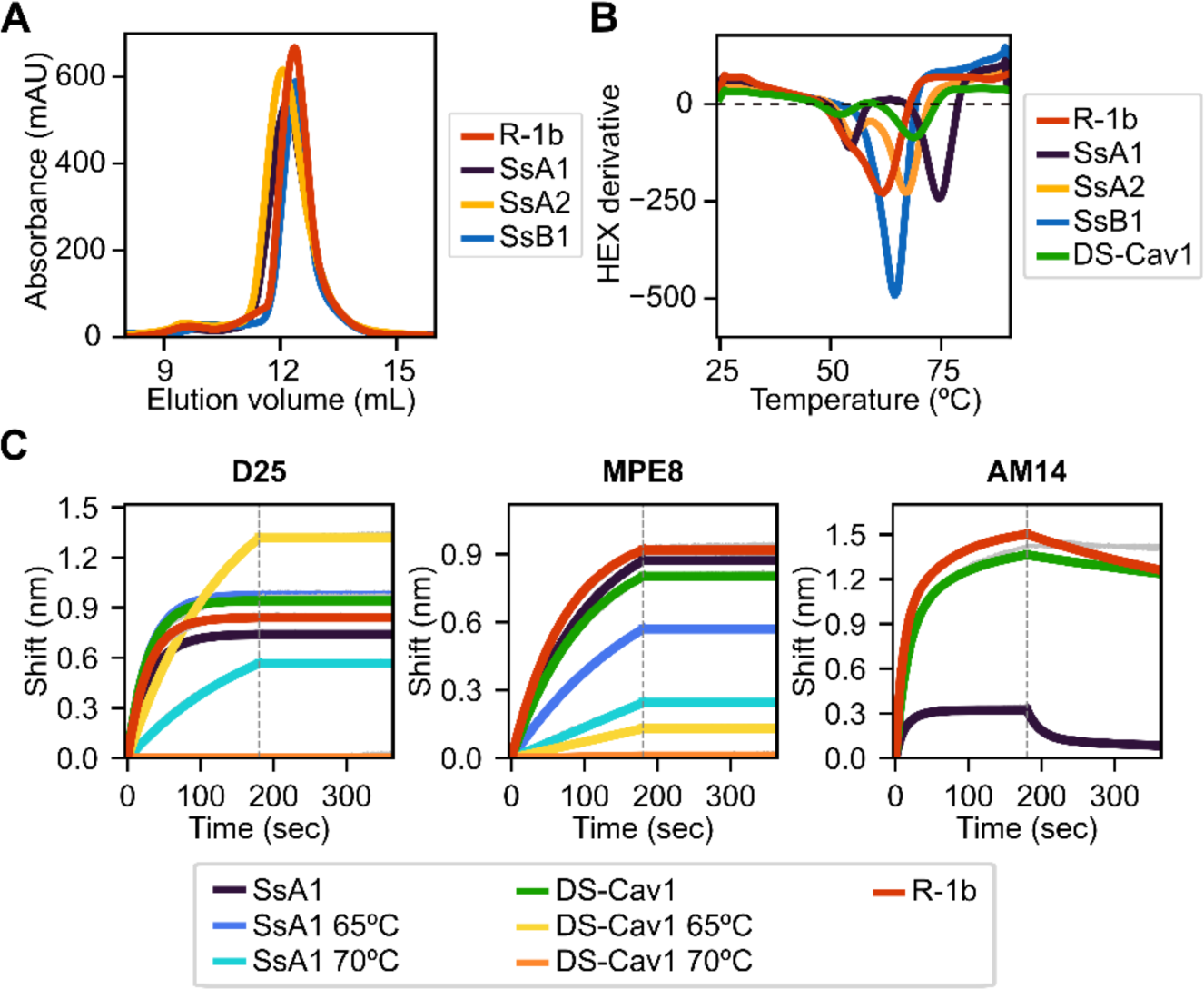
Biochemical characterization of R-1b variants containing designed non-native disulfide bonds. (A) Size-exclusion chromatography. **(B)** Differential scanning fluorimetry of designed variants in comparison with the clinical candidate DS-Cav1(7). On display are shown averaged values over three replicates. Estimated melting temperatures are R-1b 62°C, SsA1 74.5°C, SsA2 67.5°C, SsB1 64.5°C, and DS-Cav1 68°C. **(C)** Binding of variants SsA1, R-1b, and DS-Cav1 to prefusion antibodies after heat treatment. R-1b was only tested at room temperature as the protein’s melting temperature was below the set temperature for the assays. Plotted values correspond to binding at the highest protein concentration (200nM). BLI row data is shown in grey, while fitted curves are shown in colors. The end of the association time is delimited with a dotted line. Binding constants are shown in Supplementary Table 1.

The melting curve of SsA1 and DS-Cav1 presented two possible unfolding stages, characterized by a low-intensity peak at approximately 55°C and a high-intensity peak above 65°C (Fig. 3.B). To assess the structural significance of these unfolding phases, we conducted binding assays with prefusion-specific antibodies after subjecting the proteins to heat treatment. Our results revealed that SsA1 effectively preserves the prefusion conformation at high temperatures, as evidenced by its strong binding to D25 (46, 47) and MPE8 (48) even after heating at 65°C, and continued binding to D25 even after heating at 70°C (Fig. 3.C). In contrast, although DS-Cav1 retained binding to D25 at 65°C, the reduced binding to MPE8 indicated the loss of the protein’s quaternary structure at this temperature (Fig. 3.C). These results not only confirmed the increased stability of SsA1 compared to DS-Cav1 but also demonstrated that the apparent unfolding observed at ∼55°C has no substantial impact on the conformation of prefusion antigenic regions. Instead, it is the second melting peak that determines the protein’s complete unfolding. Our antibody binding assays also reveal a diminished interaction between SsA1 and AM14 (47, 49) at room temperature, indicating a potential disruption of the antigenic site V (Fig. 3.C).

### Disulfide bond detection in SsA1

To verify the successful formation of the V157C-N183C disulfide bond in design SsA1, we conducted a tandem liquid chromatography mass spectrometry (LC-MS/MS) analysis. To achieve this, the protein underwent two consecutive alkylation reactions, enabling the differentiation of free cysteines from those involved in disulfide bonding. In the first alkylation step, we used iodoacetic acid (IAA) to attach carboxymethyl groups specifically on free cysteines. Subsequently, a second alkylation reaction was carried out with iodoacetamide (IAM) to introduce carbamidomethyl groups on disulfide-bonding cysteines, following the reduction of the disulfide bond. Finally, peptide fragments obtained from trypsin digestion were subjected to LC-MS/MS analysis.

The ion chromatograms of peptides containing cysteines 157 (**<u>C</u>**LHLEGEVNK) and 183 (AVVSLS**<u>C</u>**GVSVLTSK) displayed two distinct retention times, indicating the presence of both carboxymethyl and carbamidomethyl in the peptides (Supplementary Fig. 1). Although this signified that the cysteines 157 and 183 existed in both free and bound states, the predominance of disulfide-bonding cysteines was evident in base peak intensity of the samples’ mass spectrum (Supplementary Fig. 1). Specifically, the normalization level (NL) value of each peak indicated that the relative concentration of the peptide **<u>C</u>**LHLEGEVNK in the disulfide-bonding state was ∼29 times higher than in the unbound state. Similarly, the concentration of the peptide AVVSLS**<u>C</u>**GVSVLTSK in the disulfide-bonding state was ∼15 times higher than in the unbound state. The fragmentation spectrum of both peptides, confirming the disulfide formation (carbamidomethyl labeling), is shown in Supplementary Figs. 2 and 3.

### Immunogenicity of SsA1 in cotton rats

The SsA1 protein was selected for a vaccination study to investigate the potential effects of enhanced prefusion-stability on the immunogenicity of R-1b. Cotton rats were immunized intramuscularly with 10 or 100 µg of AddaSO3-adjuvanted R-1b or SsA1 at weeks 0 and 4 (Fig. 4.A). Three weeks after the booster immunization, sera were collected to examine the IgG response against prefusion and postfusion RSV F proteins, along with their neutralizing activity (Fig. 4.A). Notably, animals immunized with either R-1b or SsA1 presented a robust RSV F prefusion-specific response, showing equivalent antibody titers against RSV A or RSV B F proteins, as measured by ELISA (Fig. 4.B and D). Similarly, no significant differences were observed in terms of RSV A2 F postfusion-specific antibodies, although SsA1 vaccination evidenced slightly lower titers (Fig. 4.C). Finally, both immunogens induced comparable levels of antibody neutralizing titers, with most doses surpassing the neutralizing titers generated by natural RSV A2 infection (Fig. 4.E).

**Figure 4.**
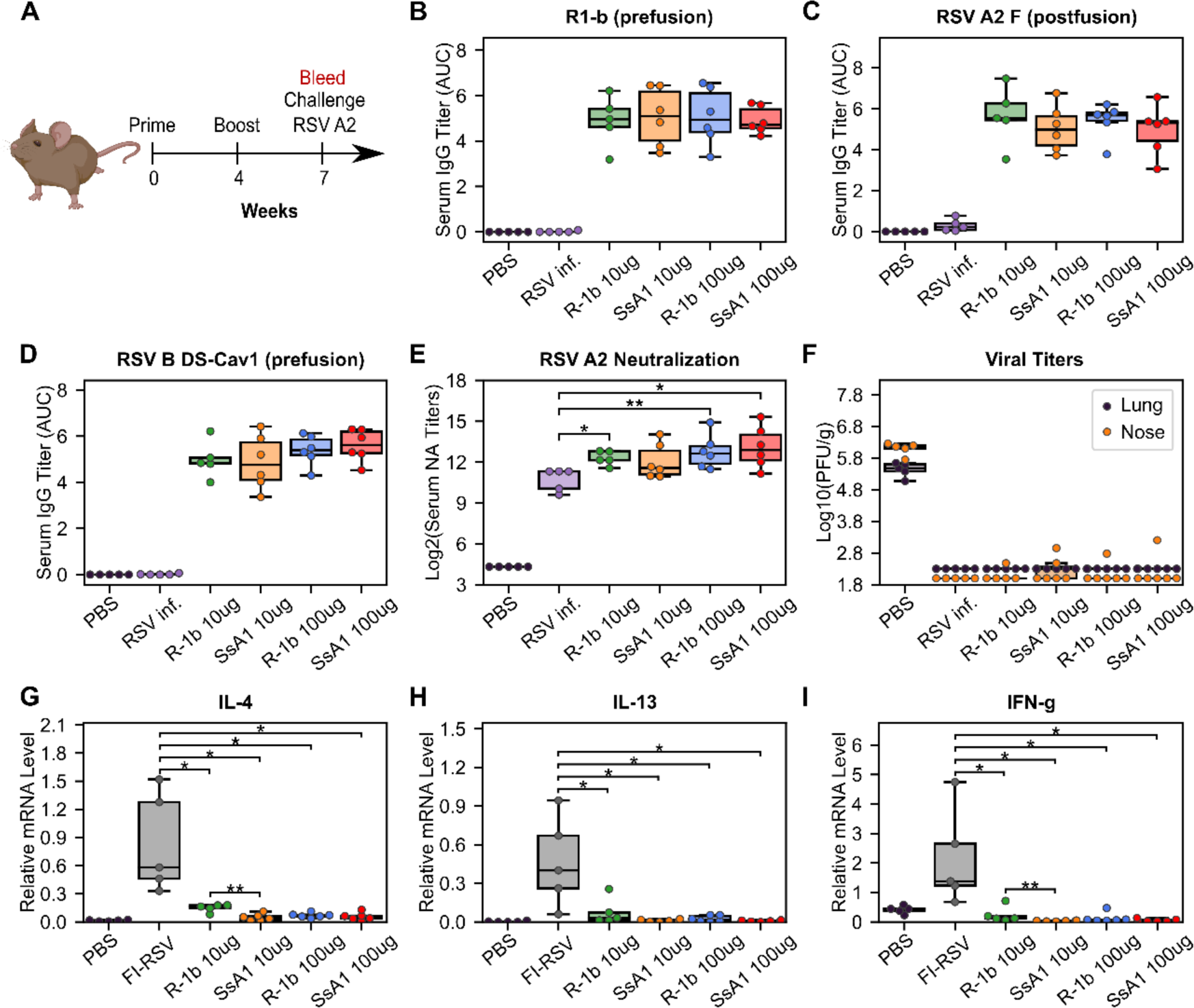
Immunogenicity and vaccine-enhanced disease assessment of R1-b variants in cotton rats. (A) Vaccination study regimen. **(B)** Serum IgG binding against R1-b, **(C)** RSV A/A2 F (postfusion) (50), and **(D)** RSV B/18537 DS-Cav1(51) proteins measured by ELISA three weeks post-boost. Control experiments corresponded to a mock vaccination with PBS and a primary infection with RSV A/A2 on day 0 (RSV inf.). Serum titers were estimated with GraphPad Prism 9.0(52)(52) using the area under the curve (AUC) measurement with a baseline of 0.3 absorbance units and log_3_-transformed serum dilutions. Sera were serially diluted three-fold from 1:60,000 (R1-b binding) or 1:20,000 to a total of 7 dilutions. **(E**) Serum neutralizing (NA) titers against RSV A/A2 using sera three weeks post-boost. **(F)** RSV A/A2 viral titers in lung and nose five days after challenge. Viral titers were determined by plaque forming units (PFU) per gram of tissue. **(G)** Vaccine-enhanced disease assessment according to mRNA levels of Interleukin-4 (IL-4)**, (H)** Interleukin-13 (IL-13), and **(I)** Interferon gamma (IFN-g). mRNA levels were measured in lung tissue five days post-challenge. A mock vaccination with PBS was used as the negative control, and cotton rats immunized with formalin-inactivated RSV (FI-RSV) were used as the positive control. The relative expression units were normalized to the level of β-actin mRNA (“housekeeping gene”) expressed in the corresponding sample. In all panels, each animal is represented by a circle corresponding to the averaged value from two measurements. The distribution of the data is shown in boxplots. All boxplots show the median as a central line, lower and upper quartiles as the box limits, and minimum to maximum values as whiskers. Pairwise statistical analyses were performed with a two-tailed T-test for the ELISA experiments and the vaccine-enhanced disease assessment. A Mann-Whitney U test was used for the neutralization data. *, p ≤ 0.05; **, p ≤ 0.01. Cotton rat cartoon was created with BioRender.

Our vaccination study concluded with an RSV A2 challenge to assess the efficacy and safety of our proteins as potential vaccine candidates. Overall, animals vaccinated with either R-1b or SsA1 successfully cleared the virus from the lung and nose (Fig. 4.F) and did not exhibit vaccine-enhanced disease, as evidenced by low levels of interleukin (IL) 4, IL-13, and interferon-gamma (IFN-g) (Fig. 4.G-I). Vaccine safety was further supported by the absence of enhanced pulmonary histopathology. Animals vaccinated with 10 µg of R-1b or SsA1 presented minimal lung lesions, which did not exceed those observed in animals mock-immunized with PBS, while no histopathology was detected in animals immunized with 100 µg doses (Supplementary Fig. 4). Collectively, our vaccination data suggests that the increased physical stability of SsA1 did not result in strengthened protection against RSV. However, it remains unclear whether the disruption of the antigenic site V hindered such potential improvement.

### Epitope-specific antibody response

To gain further insights into the epitope-specific response elicited by SsA1 and R-1b, we conducted antibody binding competition assays against D25, MPE8, and 131-2A (53). This competition was carried out following the interaction of the R-1b antigen with different concentrations of pooled serum three weeks post-boost vaccination. Consistent with our ELISA findings, the sera from SsA1 or R-1b vaccinated animals exhibited strong competition for binding sites specific to the prefusion state (Fig. 5). Notably, vaccinations with SsA1 demonstrated a higher prevalence of antibodies targeting the antigenic site Ø. In contrast, antibodies specific to the postfusion state were predominantly observed in animals vaccinated with R-1b, while quaternary antibodies were equally abundant in both vaccination groups (Fig. 5). Altogether, our results suggest that even though the overall protection against RSV seems comparable with both immunogens, there are variations in the specificity of the antibody response originating from the differences in the physical stability of their respective binding sites. Consequently, it is likely that more pronounced differences in protection may become apparent at reduced vaccination dosages.

**Figure 5.**
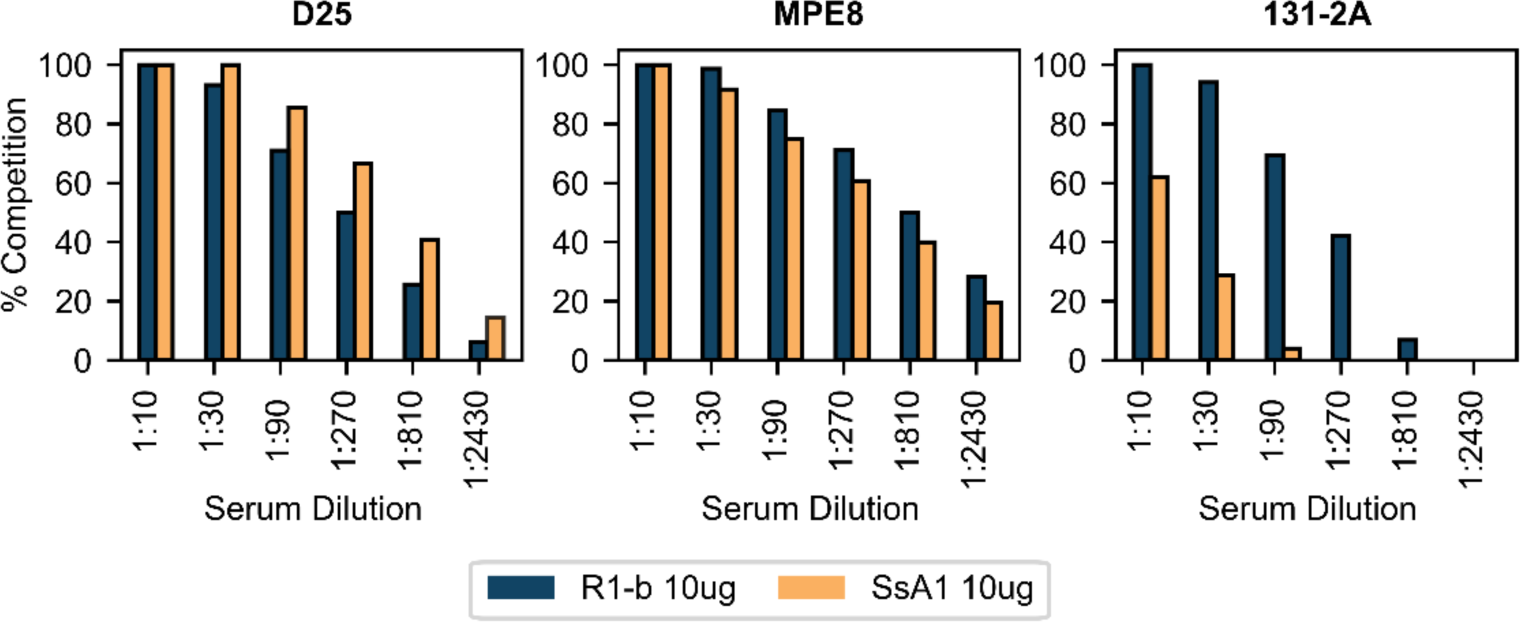
BLI-based antibody binding competition against three weeks post-boost serum. Pooled serum samples from rats vaccinated with 10 µg of R-1b or SsA1 were tested for binding competition against D25, MPE8, and 131-2A antibodies. Sera reactivity was first evaluated against immobilized R-1b and the complexes R-1b+serum antibodies were then tested for interaction with the reference antibodies. Percentage of antibody competition was calculated as 1-(association response vaccinated serum / association response naive serum) × 100%. Measurements were performed by duplicates and the averaged value is on display.

## Discussion

Given the clinical significance of prefusion class I fusion proteins (54–57), and the proven effectiveness of non-native disulfides in enhancing protein stability (1, 3, 7, 29), we have developed a computational approach to identify prefusion-stabilizing disulfide bonds. While we had previously pioneered an automated method aimed at increasing the stability of class I fusion proteins, we recognized the need to complement it due to its primary emphasis on optimizing non-covalent interactions. Indeed, our prefusion-stabilized RSV F protein, R-1b, emerged as an ideal candidate for exploring the influence of designed disulfide bonds, as its optimization was exclusively centered on reinforcing weak molecular forces (32). Notably, despite the extensive research on stabilizing the prefusion RVS F (7, 26, 30, 44, 45), our disulfide search has uncovered previously unexplored mutations. These findings highlight the comprehensive analysis that computational design offers, showcasing its potential benefits not only for novel optimizations but also for the enhancement of established vaccine candidates.

During the preparation of this manuscript, an independent study was published, assessing the impact of various mutations on the stability of the prefusion RSV F protein, including the introduction of new disulfide bonds (45). While some of our findings aligned, such as the identification of the A55C-L188C and S443C-S466C disulfides, with the former proving more stabilizing than the latter, other results were exclusive to each study. Of note, this independent investigation reported a newly discovered highly stabilizing disulfide, namely T103C-I148C. We noticed that this disulfide was overlooked with our approach due to its location in the fusion peptide region, which we had occluded. Our methodology, which initiates the scanning process by comparing prefusion and postfusion conformations, excluded the fusion peptide region because of its absence in the postfusion conformation. However, upon revisiting our scanning process, this time including the fusion peptide, the T103C-I148C disulfide was successfully identified (Supplementary Fig. 5).

While predicting the likelihood of disulfide formation is feasible through the analysis of the bond’s geometry (33–35), a metric to rank the stabilizing effect of a designed disulfide is not currently available. Some studies have proposed using residue B-factors to select stabilizing disulfides, recognizing that rigidifying highly flexible areas can enhance protein stability (33, 36). However, this metric may not fully capture the complexities of the prefusion state in class I fusion proteins. The RSV F protein serves as a clear example. Depending on the crystallization method and quality, the RSV F region with the highest B-factors can vary between the head and the membrane-proximal domains, as evident in the PDBs 5c69 and 5w23 (30, 58), respectively. Interestingly, despite this variability, one of the most stabilizing disulfides reported for RSV F, the S155C-S290C disulfide (7), falls in a region of relatively low B-factor. This observation underscores the challenges of predicting disulfide-induced stabilization of the prefusion conformation and emphasizes the need to consider other factors when estimating the potential success of an engineered disulfide bond.

As class I fusion proteins undergo intricate refolding processes, we explored whether the dynamics of their conformational changes could provide insight into the ability of a non-native disulfide to stabilize the prefusion conformation. To investigate this, we employed the RMSD of the prefusion-to-postfusion transition as a quantitative metric for monitoring conformational dynamics. We hypothesize that regions with high RMSD are more prone to unfolding first during the transition to the postfusion state, and inhibiting these initial rearrangements is more effective at stabilizing the prefusion conformation. Our results seem to support this hypothesis, revealing poorer thermal stability in disulfides located in regions of smaller RMSD values. However, since conformational changes in the RSV F protein probably initiate in the refolding region 1 followed by the refolding region 2 (30), further studies are essential to clarify whether the overall RMSD value can inform the stabilizing properties of a disulfide, or if its utilization requires subdivision based on refolding regions.

For the RSV F protein, we have found that disulfides placed in proximity to the fusion peptide have a greater effect on the stability of the prefusion state than those located nearer to the membrane-proximal region. Interestingly, within the refolding region 1, increases in melting temperature aligned with the positioning of the designed disulfides, with bonds closer to the N-terminal of the F1 subunit providing higher thermal stability (Fig. 3.B). These results might reflect the dynamics of the protein, suggesting that disulfide bonds restricting the initial stages of the conformational switch have a more significant impact on the stability of the prefusion state than those hindering the unfolding at later stages. However, since the precise sequence of events preceding the translocation of the fusion peptide has not been definitively determined, and it is still unclear whether the release of the fusion peptide promotes the refolding of the heptad repeat A region or if conformational changes at the apex initiate the projection of the fusion peptide (30). Our findings appear to lend support to the former hypothesis. If conformational shifts were to initiate at the apex and propagate downstream, our A55C-L188C disulfide, located closer to the apex, should have exhibited the most significant stabilizing effect by hindering the initial unfolding phase. Contrarily, it was the V157C-N183C disulfide, situated closer to the fusion peptide, which displayed the most pronounced stabilizing impact. Therefore, it is likely that the release of the fusion peptide drives the subsequent conformational changes and its inhibition results in a more pronounced stabilization of the prefusion state, as it has been seen in other viral families (59).

At this point, it remains to establish a clear correlation between the stability of the prefusion state and its immunogenic properties which could be examined by reducing the overall dosage. It is likely that the disruption of the antigenic site V in SsA1 impeded its potential for enhanced immunogenicity (45). Prior studies aiming to improve the DS-Cav1 stability have shown a relationship between physical stability and increased RSV protection, particularly when employing interprotomer disulfides to ensure an appropriate quaternary structure (26). However, this association is not consistently observed, as other studies incorporating interprotomer disulfides and increased stability have not reported a corresponding increase in immunogenicity (44, 60). Our observations from previously reported studies (26, 44, 45, 60) indicate that disulfides involved in enhanced immunogenicity are primarily stabilizing antigenic regions. Hence, it is probable the local stability of antigenic regions might be the determining factor to increase immunogenicity rather than the overall protein stability. Within our study, we did notice that antibodies targeting the antigenic site Ø, which was stabilized by the disulfide V157C-N183C, were slightly higher in vaccinations with SsA1 (Fig. 5). However, this effect could also be attributed to the disruption of the antigenic site V and a potential reduction in interclonal competition. Nevertheless, despite the complexities surrounding the relationship between stability and immunogenicity, strategic placement of disulfide bonds may offer opportunities to enhance the quality of the antibody response.

## Materials and Methods

### I. Computational approach to design disulfide bonds

All computational analyses were performed with the Rosetta version: 2020.10.post.dev+12.master.c7b9c3e c7b9c3e4aeb1febab211d63da2914b119622e69b

#### Definition of conformational switch area

To ensure the RSV F protein is locked in its prefusion state, we focused on restraining the mobility of regions undergoing drastic conformational changes. These flexible areas were automatically identified by calculating the root-mean-square-deviation (RMSD) of each Cα atom in the prefusion conformation (R-1b protein, PDB: 7tn1) (32) compared to the postfusion structure (PDB: 3rrt) (50). Residue positions displaying motion levels of at least 10Å were considered to be involved in the conformational switch and selected for disulfide scanning.

#### Structure preparation

Recognizing the dynamic nature of protein structures, we generated an ensemble of conformations for the R-1b protein to conduct a comprehensive exploration of new disulfide bonds. This methodology is especially pertinent for disulfide scanning since the identification of potential disulfides is significantly influenced by the distances between atoms. A total of 200 possible structures of the R-1b protein were produced by refining its crystal structure (PDB: 7tn1) with the Rosetta relax application (39–41). This refinement process was guided by electron density data to prevent substantial deviations from the initial protein configuration (61). The protein’s density map was reconstructed from its map coefficients using the Phenix software version 1.15 (62) and the option “create map from map coefficients” (region padding= 0, and grid resolution factor= 0.3333). The relaxation process was carried out by assigning a weight of 35 to the density energy term of the Rosetta scoring function and performing five rounds of rotamer packing and minimization (39).

#### Design of disulfide bonds

For each conformation within the protein ensemble and every residue within the conformational switch area, we identified pairs of residues to be mutated to cysteine based on a Cβ-Cβ distance ≤ 6Å. Subsequently, disulfide formation was enforced using the Rosetta Disulfidize mover and the PyRosetta interface (37, 38). Disulfides displaying rotation/translation distances “match_rt_limit” < 2 and disulfide scores “dslf_fa13” < 3.5 were selected for further refinement. This refinement was executed using the Rosetta FastRelax mover in RosettaScripts (39, 40, 63, 64). During the relaxation process, backbone energy minimization was permitted in segments of 10 residues up and downstream of the disulfide pair, while protein repacking was confined to a 10Å radius around the new bond. A total of five rounds of relaxation were conducted.

#### Selection of candidate disulfide bonds

To minimize significant disruption to the overall protein structure, disulfide designs were initially filtered based on a total RMSD value < 0.5Å compared to their parent configuration. Furthermore, we eliminated any non-native disulfides within a 7Å distance from native disulfides to prevent nonspecific interactions. Subsequently, to increase the likelihood of successful disulfide formation, we implemented a two-step filtering process to ensure the adequacy of the disulfide geometry. At first, we screened all predicted disulfides using the Rosetta disulfide geometry potential “dslf_fa13” with a threshold of < 0.1. Following this, we compared our disulfide bond angles (dihedrals χ1, χ2, χ3, and angle Cβ-S-S) to a dataset of 300 high-resolution disulfide-containing structures (<1.5 Å) sourced from the Protein Data Bank. This dataset underwent prior relaxation using RosettaScripts and the FastRelax mover. Our disulfide candidates were then filtered based on their alignment with the observed angle ranges in these reference proteins.

In addition to evaluating the geometry, we ensured that newly introduced disulfides did not disrupt protein packing, especially when cysteine substitution involved buried hydrophobic residues. To achieve this, we identified buried hydrophobic residues in the monomeric parent structure using PyRosetta and considering a solvent-accessible surface area (SASA) ≤ 20Å² with a 2.2 Å probe size (65). Next, we assessed the packing quality within a 7Å radius around each buried residue using the PackstatCalculator (66) over five repetitions. This calculation was conducted both in the disulfide-containing model and the parent structure. Disulfide candidates were then considered when the packing quality was not disturbed by more than 0.2 units compared to the parent structure.

Finally, we incorporated the conformational dynamics of the RSV F protein to rank the potential prefusion-stabilization effect of the introduced disulfides. For this purpose, we utilized the root-mean-square deviation (RMSD) of the prefusion-to-postfusion transition as a quantitative metric to evaluate conformational dynamics. Our hypothesis suggests that regions with high RMSD are more likely to unfold first during the transition to the postfusion state. Therefore, disulfides hindering these initial rearrangements would be more effective at stabilizing the prefusion state. As a result, we ranked our disulfide candidates based on the highest RMSD observed within the residue pair forming the bond. This ranking was performed independently for the refolding region 1 and 2, as the conformational changes in the refolding region 2 depend on the earlier changes in refolding region 1 (30). The RMSD values were calculated using the same methodology employed for the identification of the conformational switch area.

### II. Protein expression and characterization

#### Protein expression

Three R-1b disulfide-containing variants, namely SsA1 (V157C /N183C), SsA2 (A55C/L188C), and SsB1 (S443C/S466C), and the control R-1b, RSV A/A2 F (postfusion) (50), RSV A/A2 DS-Cav1 (7), and RSV B/ 18537 DS-Cav1 (51) constructs were expressed by transient transfection of FreeStyle 293-F cells (Thermo Fisher) with polyethylenimine (PEI) (Polysciences). All R-1b variants were produced in pCAGGS plasmids encoding the R-1b protein, a C-terminal T4 fibritin trimerization motif (Foldon), and a His6-tag. The sequence of the R-1b protein contained residues 1-105 and 137-513 with the respective substitutions under study, and a short linker replacing the furin cleavage site and the p27 peptide (“QARGSGSGR”)(30). DNA sequences were codon optimized for human expression using the online tool GenSmart Codon Optimization (67). 293-F cells were incubated at 37°C and 8% CO_2_ for three days after transfection, and proteins were purified by nickel affinity chromatography followed by size-exclusion chromatography (SEC). SEC was carried out using a Superdex200 column (Cytiva) and phosphate-buffered saline (PBS) buffer pH 7.4.

#### Thermal stability

Differential scanning fluorimetry (DSF) was used to monitor protein stability as a function of temperature. The samples analyzed contained 4µM of protein, 5X SYPRO orange fluorescent dye (Thermo Fisher), 5 mM MgCl_2_, 50 mM KCl, and 50 mM Tris (pH 7.4). All measurements were performed by triplicates using a qPCR instrument (CFX Connect, BioRad) and a temperature gradient from 25 to 90°C with 0.5°C increments. The melting temperature of each protein was estimated based on the lowest point of the negative first derivative of the fluorescence signal.

#### Antigenic characterization

Antigenic preservation of prefusion epitopes was evaluated by bio-layer interferometry (BLI) using the prefusion-specific antibodies D25 (46, 47) (Thermo Fisher), AM14 (47, 49) (Cambridge Biologics), and MPE8 (48). Binding against expressed designs was tested with 15 nM of the antibodies and eight concentrations of the antigens, starting from 200 nM and decreasing by two-fold dilutions. Prior to all binding assays, Protein A biosensors (GatorBio) were equilibrated for 20 min in BLI buffer (PBS buffer supplemented with 0.02% tween-20 (Promega) and 0.1% bovine serum albumin (BSA) (Sigma)). Subsequently, immobilization of the antibodies on the biosensor tips was allowed for 180s, followed by a baseline correction of 120s, and an association and dissociation steps of 180s each. All assays were performed on a GatorPrime BLI instrument (GatorBio) at a temperature of 30°C and frequency of 10 Hz. Binding constants were obtained with the GatorOne software 1.7.28, using a global association model 1:1 for D25 and MPE8, and 2:1 for AM14.

Antigenic preservation after heat treatment was evaluated in the R1-b variant with the highest melting temperature (SsA1) and the control proteins R1-b and RSV A/A2 DS-Cav1. The proteins were incubated for one hour at 65 and 70°C in a thermocycler with heated lid (T100, BioRad) Binding to D25 and MPE8 antibodies was measured afterwards following the protocol described above.

#### Disulfide bond detection through alkylation and mass spectrometry

Alkylation with iodoacetic acid (IAA) and iodoacetamide (IAM) was used to corroborate the formation of the disulfide bond V157C - N183C in the SsA1 protein. These alkylation reactions were intended to label free cysteines differentially from disulfide-bonding cysteines. Specifically, carboxymethyl groups were attached to free cysteines (IAA reaction) and carbamidomethyl groups were attached to disulfide-bonding cysteines (IAM reaction), after reduction of the disulfide bond. Following alkylation, the samples were digested, and peptides were analyzed through tandem liquid chromatography mass spectrometry (LC-MS/MS).

The detailed steps of the process are described below:

- *Alkylation reaction:* 20 µL of SsA1 protein at 1.5 mg/mL were incubated for 30 minutes in the dark with 2 µL of 0.1M iodoacetic acid. The protein sample was denatured and reduced by adding 0.5 µL of 0.5M dithiothreitol (DTT). The mixture was heated at 100°C for 5 minutes and after cooling down the reaction was allowed for 30 minutes at room temperature. The second alkylation was carried out by adding 2 µL of 0.5M iodoacetamide and incubating for 30 minutes.
- *Deglycosylation:* The alkylated sample was mixed with 100 µL of digest buffer (0.2% sodium deoxycholate (SDC) in 50 mM ammonium bicarbonate) and digested with 20U PNGase F (Lectenz Bio, Athens) in Sartorius Vivacon 500 (10K MWCO) for 2.5 hours at 37°C.
- *Trypsin digestion:* The sample in the filter was washed twice with 200 µL of 20 mM triethylammonium bicarbonate. 0.3 µg of Trypsin in 50 µL of 20 mM triethylammonium bicarbonate were added to the sample in the filter, and the digestion was carried out overnight at 37°C. The next day, the tryptic digests were spun out of the filter, and remaining peptides in the filter were eluted with 100 µL of water. The tryptic peptides were dried by Vacufuge.
- *LC-MS/MS:* The mass spectrometry analyses were performed on a Thermo Fisher LTQ Orbitrap Elite Mass Spectrometer coupled with a Proxeon Easy NanoLC system (Waltham, MA) located at Proteomics and Mass Spectrometry Facility, University of Georgia. The enzymatic peptides were loaded into a reversed-phase column (self-packed column/emitter with Dr. Maisch ReproSil-pur C18AQ 120Å 3 μM resin), and directly eluted into the mass spectrometer. Briefly, the two-buffer gradient elution (0.1% formic acid as buffer A and 99.9% acetonitrile with 0.1% formic acid as buffer B) started with 0% buffer B for 2 minutes, and then increased to 40% buffer B for 95 minutes and to 95% buffer B for 10 minutes. The MS data was obtained using the Xcalibur software (version 3.0, Thermo Fisher Scientific) and the data-dependent acquisition (DDA) method. A survey MS scan was acquired first, and then the top 10 ions in the MS scan were selected for following CID (collision-induced dissociation) and HCD (higher energy C trap dissociation) tandem mass spectrometry (MS/MS) analysis. Both MS and MS/MS scans were obtained by Orbitrap at the resolutions of 120,000 and 15,000, respectively. Protein identification and modification characterization were performed using Thermo Proteome Discoverer (version 3.0) with Mascot (Matrix Science) against Uniprot plus the SsA1 sequence, and a modified contaminations database with commonly known contaminating proteins (Mascot).

### III. Animal studies

#### Cotton rat immunization

Inbred 6-8 weeks-old, *Sigmodon hispidus* female and male cotton rats (source: Sigmovir Biosystems, Inc., Rockville MD) were maintained and handled under veterinary supervision in accordance with the National Institutes of Health guidelines and Sigmovir Institutional Animal Care and Use Committee’s approved animal study protocol (IACUC Protocol #15). Cotton rats were housed in clear polycarbonate cages and provided with standard rodent chow (Harlan #7004) and tap water *ad lib.* Groups of 5 or 6 animals (3 females/2 males or 3 females/3 males) were immunized intramuscularly with two different doses (10 µg or 100 µg) of either purified R-1b or SsA1 protein with AddaSO3 adjuvant (50% v/v) at weeks 0 and 4 (Prime and Boost) (Fig. 4.A). Control experiments were carried out by immunizing animals only with PBS (negative control), inoculating animals intranasally with 10^5^ PFU of RSV A/A2 Live on week 0 (positive control for RSV A/A2 infection), or by immunizing animals with formalin-inactivated RSV (FI-RSV lot#100) diluted 1:100 in PBS (positive control for vaccine-enhanced disease). Bleeds were collected from the retro-orbital sinus at week 7 (Fig. 4.A), and sera were analyzed by ELISA, BLI, and neutralization assay.

#### RSV A/A2 challenge

RSV A/A2 (ATCC, Manassas, VA) was propagated in HEp-2 cells after serial plaque-purification to reduce defective-interfering particles. A pool of virus designated as RSV/A2 Lot# 092215 SSM containing approximately 3.0 x 10^8^ pfu/mL in sucrose stabilizing media was used for the *in vivo* experiment. Virus stock was stored at −80°C and had been characterized *in vivo* using the cotton rat model and validated for upper and lower respiratory tract replication. Vaccinated and control animals were inoculated intranasally at week seven with 0.1 mL of RSV/A2 (Lot# 092215 SSM) at 10^5^ PFU per animal. Back titration on the challenge virus was performed to confirm challenge dose. Cotton rats were euthanized on day five post-infection for analysis of viral load and lung histopathology and mRNA gene expression.

#### RSV IgG measurement by ELISA

One hundred (100) µL of purified R-1b (32), RSV A/A2 F (postfusion) (50), RSV A/A2 DS-Cav1 (7), and RSV B/ 18537 DS-Cav1 (51) at 2 µg/mL were coated onto 96 well ELISA plates (Immulon 2 HB, Thermo Fisher) at 4°C overnight. Next day, plates were washed with wash buffer (PBS buffer with 0.05% tween-20) and blocked with 200 µL/well of blocking buffer (wash buffer supplemented with 3% non-fat milk (LabScientific) and 0.5% BSA). After two hours of incubation at room temperature, plates were washed, and 100 µL/well of diluted serum from each rat was added and incubated for two hours at room temperature. Serum was serially diluted 1:3 in blocking buffer starting from a 1:20,000 or 1:60,000 (R1-b binding) dilution until a total of seven dilutions. Once the serum incubation finished, the plates were washed, and 100 µL/well of Rabbit anti Cotton Rat IgG (1:1,000 in blocking buffer) (Invitrogen) was added and incubated for one hour at room temperature. The plates were washed again and 100 µL/well of Goat anti Rabbit IgG-HRP (1:9,000 in blocking buffer) (Invitrogen) was added and incubated for one hour at room temperature. Finally, plates were washed and 100 µL/well of TMB substrate solution (Fisher Scientific) was added and incubated in the dark for 15 minutes. The reaction was stopped with 100 µL/well of 2M H_2_SO_4_ and the optical density (OD) was measured at 450 nm on a SpectraMax M2 Reader (Molecular Devices). All experiments were performed by duplicates and the averaged OD value was considered for analysis. Serum titers were estimated with GraphPad Prism 9.0 (52) using the area under the curve (AUC) measurement with a baseline of 0.3 absorbance units and log_3_-transformed serum dilutions.

#### Antibody competition assays using BLI

Antibody binding competition against D25, MPE8 and 131-2A (53) (Millipore Sigma) was carried out to estimate epitope-specific responses after vaccination. Prior to data collection, anti-penta-His sensors (GatorBio) were hydrated for 20 minutes in BLI buffer (PBS buffer supplemented with 0.5% BSA and 0.05% tween-20, pH 7.4). The assay was initiated with an equilibration step in BLI buffer for 100s followed by immobilization of the R-1b or RSV A2 F (postfusion) proteins at 20 µg/mL, for 50s. The probes were then washed for 200s in blocking buffer (25% ChonBlock buffer (Chondrex Inc.) diluted in BLI buffer) and interaction with pooled serum samples (three weeks post-boost) was allowed for 700s. Sera were serially diluted 1:3 in blocking buffer, starting from a 1:10 dilution until a total of six dilutions. Blocking buffer with no serum and naïve serum from PBS-vaccinated rats (1:10 dilution) were also included as controls. After serum interaction, a baseline phase was carried out by dipping the probes in blocking buffer for 60s. Finally, association of competing antibodies was performed for 700s. The MPE8 antibody was tested at 18 µg/mL while the remaining antibodies were tested at 9 µg/mL. All antibodies were diluted in blocking buffer and the antigens were diluted in BLI buffer. Percentage of antibody competition was calculated as 1-(association response vaccinated serum / association response naïve serum) × 100%. Measurements were performed by duplicates and the averaged competition value was considered for analysis. All assays were performed with a GatorPrime BLI instrument at a temperature of 30°C and frequency of 10 Hz.

#### RSV neutralizing antibody assay (60% reduction)

Heat inactivated serum samples were diluted 1:20 with Eagle’s Minimum Essential Medium (EMEM) and serially diluted further 1:4. Diluted serum samples were incubated with RSV A/A2 (25-50 PFU) for one hour at room temperature and inoculated in duplicates onto confluent HEp-2 monolayers in 24 well plates. After one hour incubation at 37°C in a 5% CO2 incubator, the wells were overlayed with 0.75% methylcellulose medium. After four days of incubation, the overlay was removed, and the cells were fixed with 0.1% crystal violet stain for one hour and then rinsed and air dried. The corresponding reciprocal neutralizing antibody titers were determined at the 60% reduction end-point of the virus control. The averaged value for two measurements was considered for analysis.

#### Lung and nose viral titration

Lung (*en bloc* and tri-sect, left section) and nasal tissue were homogenized in 3 mL of Hanks’ Balanced Salt Solution (HBSS) supplemented with 10% Sucrose-Phosphate-Glutamate (SPG). Homogenates were clarified by centrifugation and diluted in EMEM. Confluent HEp-2 monolayers were infected in duplicates with diluted homogenates in 24 well plates. After one hour incubation at 37°C in a 5% CO_2_ incubator, the wells were overlaid with 0.75% methylcellulose medium. After 4 days of incubation, the overlay was removed, and the cells were fixed with 0.1% crystal violet stain for one hour and then rinsed and air dried. Plaques were counted and virus titer was expressed as plaque forming units per gram of tissue. The averaged value over two measurements was used for analysis.

#### Real-time PCR

Total RNA was extracted from homogenized tissue (lingular lobe) or cells using the RNeasy purification kit (QIAGEN). One µg of total RNA was used to prepare cDNA using Super Script II RT (Invitrogen) and oligo dT primer (1 µL, Invitrogen). For the real-time PCR reactions, the Bio-Rad iQ^TM^ SYBR Green Supermix was used in a final volume of 25 µL, with final primer concentrations of 0.5 µM. Reactions were set up in duplicates in 96-well trays. Amplifications were performed on a Bio-Rad iCycler for 1 cycle of 95°C for 3 min, followed by 40 cycles of 95°C for 10s, 60°C for 10s, and 72°C for 15s. The baseline cycles and cycle threshold (Ct) were calculated by the iQ5 software in the PCR Base Line Subtracted Curve Fit mode. Relative quantitation of DNA was applied to all samples. The standard curves were developed using serially-diluted cDNA sample most enriched in the transcript of interest (*e.g.*, lungs from 6 hours post RSV infection of FI-RSV-immunized animals). The Ct values were plotted against Log_10_ cDNA dilution factor, and these curves were used to convert the Ct values obtained for different samples to relative expression units. The averaged relative expression units for two measurements were then normalized to the level of β-actin mRNA (“housekeeping gene”) expressed in the corresponding sample.

#### Pulmonary histopathology

The right section of the lungs was dissected and inflated with 10% neutral buffered formalin to their normal volume, and then immersed in the same fixative solution. Following fixation, the lungs were embedded in paraffin, sectioned, and stained with hematoxylin and eosin (H&E). Four parameters of pulmonary inflammation were evaluated: peribronchiolitis (inflammatory cell infiltration around the bronchioles), perivasculitis (inflammatory cell infiltration around the small blood vessels), interstitial pneumonia (inflammatory cell infiltration and thickening of alveolar walls), and alveolitis (cells within the alveolar spaces). Slides were scored blind on a 0-4 severity scale. The scores were subsequently converted to a 0 −100% histopathology scale.

#### Statistical analysis

All statistical analysis were performed in R studio 4.1.0 using the tidyverse and rstatix packages. ELISA results and vaccine-enhanced disease assessment from R-1b and SsA1 vaccinations were compared applying a two-tailed t-test. Neutralization and histopathology data was compared with the Mann-Whitney U test. The normality of each group was confirmed by a Shapiro wilk test, and the equality of variances was checked through the Levene’s test. The overall level of significance was set at 5%. Boxplot visualization was done with the seaborn package in python 3.7.

## Acknowledgments

We would like to thank Sonnie Kim (NIH/NIAID) for managing the preclinical testing program and the Proteomics and Mass Spectrometry Facility at the University of Georgia, particularly Dr. Chau-wen Chou, for the mass spectrometry analysis of our samples. We would like to thank Luki Goldschmidt for his assistance in maintaining the computing resources, and Dr. Jarrod Mousa for discussions. This work was supported by federal funds from the National Institutes of Health grant R01AI140245 (KJG, EMS), National Institutes of Health Contract Number HHSN272201700028I/75N93023F00001 (KY, JB, MSB). Orbitrap was purchased with National Institutes of Health grant S10RR028859 to Professor I Jonathan Amster at the University of Georgia.

## Data and Materials Availability

SsA1 plasmid is available from EMS under a material transfer agreement with the University of Georgia. Rosetta is available through licensing https://www.rosettacommons.org. Scripts for generating designs will be available on https://github.com/strauchlab/disulfide_design upon publication.

## Author Contributions

Conceptualization: KJG, EMS; Methodology: KJG, KY, JB, MSB, EMS; Formal analysis: KJG, KY, JB, MSB, EMS; Writing–original draft: KJG, EMS; Writing–reviewing & editing: KJG, MSB, EMS; Visualization: KJG, EMS; Supervision: EMS; Project administration: EMS; Funding acquisition: EMS.

## Classification

Biological Sciences, Computational Biology.

